# Fluid shear stress coupled with narrow constrictions induce cell type-dependent morphological and molecular changes in circulating tumor cells

**DOI:** 10.1101/722306

**Authors:** Hamizah Ahmad Cognart, Jean-Louis Viovy, Catherine Villard

## Abstract

Cancer mortality mainly arises from metastases, due to cells that escape from a primary tumor, circulate in the blood as circulating tumor cells (CTCs), permeate across blood vessels and nest in distant organs. It is still unclear how CTCs overcome the harsh conditions of fluid shear stress and mechanical constraints within the microcirculation. Here, a model of the blood microcirculation was established through the fabrication of microfluidic channels comprising constrictions. Metastatic breast cancer cells of epithelial-like and mesenchymal-like phenotypes were flowed into the microfluidic device. These cells were visualized during circulation, analyzed for their dynamical behavior and retrieved post-circulation. γ-H2AX staining showed significant increase of DNA damage response in epithelial-like SK-BR-3 cells, while gene expression analysis of key regulators of epithelial-to-mesenchymal transition revealed significant increase of Twist2 relative expression in mesenchymal-like MDA-MB-231 cells post-circulation. This work documents first results of the changes at the cellular, subcellular and molecular scales induced by the two main mechanical stimuli arising from circulatory conditions.

## Introduction

Understanding the process of metastasis is a major challenge in the fight against cancer. This process is a multi-step one that often involves cells migrating from a primary tumor (at 90% of epithelial origin, i.e. carcinomas) into the blood stream (intravasation), where they reach a distant organ by re-crossing the endothelial barrier (extravasation). At both cellular and molecular levels, and in most cases, these events involve the ability of cells to undergo complex reprogramming processes named the epithelial-to-mesenchymal transition (EMT) at the primary tumor site, followed by the reverse process called MET (mesenchymal-to-epithelial transition), which may help in establishing distant micro-metastases after the circulation step.^1–4^ Blood circulation, together with gene expression reprogramming, therefore plays a central role in the metastatic cascade.

Once in the bloodstream, circulating tumor cells (CTCs) experience shear stress. In addition, considering that capillaries do not have uniform diameters but are regularly constricted instead^5^, CTCs encounter various constraints within the microvascular network, leading to repeated mechanical deformation and eventually to arrest. Although numerous studies have revealed important influences of mechanical constraints in cancer^6^, surprisingly only a few works explored the responses of cancer cells to the mechanical stressors provided by the blood circulation, and most of them were limited to the study of the effect of shear stress. Malignant cells appeared more resistant than non-malignant cells to shear stress^7, 8^ yet they still underwent significant apoptosis^9^, in particular as compared to hematopoietic cells.^10^ Altered cytoskeleton organization in suspended and sheared primary ovarian cells was observed.^8^ Interestingly, the process of suspending cells in and of itself triggers myosin-II inhibition, leading to increased cell stiffness.^11^ All these studies, however, involved flow-induced shear without topographical constraints, and relied on macroscopic tools that limited the control of the cell treatment uniformity, preventing single cell studies. The investigated biomarkers were also rather limited, targeting essentially cell survival.

Nonetheless, several works have used microfluidic systems to address the effect of the microvasculature geometry on cells. Pioneering work by Preira et al. investigated the role of cell deformability in the pathological arrest of leukocytes in the blood microcirculation of the lungs, by devising a microfluidic system with evenly spaced constrictions.^12^ In the field of cancer, using circulated glioblastoma and normal glial cell lines, Khan et al. demonstrated the cell entry time into the confined space provided by 11 µm-wide diameter microchannels as a better marker of malignancy than deformability.^13^ Au et al. showed that clusters of CTCs could reorganize reversibly in order to traverse microchannels of 5-10 µm-wide.^14^ Nath et al. flowed HeLa cells across 7 µm-wide constrictions, and demonstrated that cell viability was reduced by 50%, but that the expression of MMP2, a metalloproteinase involved in stromal tissue degradation, was unchanged.^15^ In a very interesting study, Xia et al. flowed leucocytes, MDA-MB-231 and MCF-7 cell lines into arrays of pores, which demonstrated a pressure dependence of the deformation of cells and nuclei, proposing that such studies could guide the optimization of CTC sorting devices.^16^ Overall, microfluidic attempts at mimicking cancer cells in the blood circulation are still sparse, and are mostly focused on a phenomenological investigation of their mechanical properties.

The question of the blood circulation effect on the cell phenotype and its genome, and ultimately aggressiveness is hardly addressed. The acquisition and maintenance of the key mesenchymal phenotype for the metastatic process, involving i.e. up-regulation of vimentin and down-regulation of E-cadherin expressions, require important cellular reprogramming by the activation of master regulators including transcription factors such as Snail, Twist and zinc-finger E-box-binding (ZEB), and transforming growth factor, TGF-β.^17, 18^ These phenotypic changes, however, are not bimodal, and recent studies suggest that disseminating tumor cells may present diverse and heterogeneous combinations of epithelial and mesenchymal phenotypic traits.^14, 19, 20^

In this article, the main aim is to tackle the least studied component of the metastatic cascade to date, which is the transient circulation step in the blood stream. More precisely, we will explore the combined influence of shear stress and physical constraints on the characteristics of circulating cancer cells. The main questions that will be addressed are how do circulation and constrictions regulate the phenotype, genome integrity and gene expression of CTCs. To address these questions, cells of two different origins, i.e. epithelial (SK-BR-3) and mesenchymal (MDA-MB-231) breast cancer cell lines, were circulated in microfluidic channels using a flow control system to recapitulate the pressure-velocity patterns of the microcirculation blood flow. While under circulation, the tumor cells were constrained due to the presence of evenly-spaced multiple constrictions to mimic the mechanical condition in the blood capillary bed. The ways in which shear flow and mechanical constraints can promote changes in gene expression were investigated and elaborated based on the common framework for EMT and its transcription factors (EMT-TFs). Our results showed that circulation affects the cells at several scales, i.e. at cellular, sub-cellular and molecular levels, and that some of these changes are modulated by the epithelial or mesenchymal initial cell type. Therefore, the role of the circulation step seems to go beyond a simple disseminating function of cancer cells to distant organs. For this reason, it should be considered as an active step that is likely to modify gene expression of CTCs and possibly their mechanical properties.

## Results

### Characterization of Metastatic Breast Cancer Cells under Circulation, Confinement and Constrictions

To test the effects of different geometric mechanical constrictions on the flow-induced migration of single tumor cells, five types of geometric microfluidic models were fabricated (Figure 1a). All five types comprised channels 420 μm in length. The “unconfined” type had a channel height of 20 μm, a value reduced to 15 µm in the “confined” type. Confined type 1, 2 and 3 comprised channels with multiple (11, 7 and 5 respectively) 6 μm-wide constrictions spaced apart by 26 μm-wide chambers that were 20 μm in length. These microfluidic designs were calculated for their flow resistance based on Newtonian fluid flow resistance (Figure 1b). A pressure set-point of 10 kPa applied across the whole microfluidic circuit was chosen in order to achieve the same order of magnitude of flow rate (i.e. approximately 1 µl/min) as the blood flow rate reported in micron-sized capillaries *in vivo*.^21^ Either poorly (SK-BR-3) or highly (MDA-MB-231) metastatic breast cancer cell lines were delivered into these five types of geometric microfluidic models for single cell mechanical phenotyping by flow-induced migration. The velocity of these cells through the micro channels was measured (Figure 1c), with values in the range of a few mm/s in agreement with recent *in vivo* values reported in the literature.^22^ More precisely, mean values close to 18 mm/s were found in the unconfined condition for both cell lines, reduced to about 10 mm/s and 8 mm/s in the confined situation for SK-BR-3 and MDA-MB-231 cells, respectively. Then, velocity data of both cell lines displayed a general trend, in which a longer constriction tends to systematically induce a lower mean velocity, with mean values of 2.8 mm/s and 5.2 mm/s in the 60 µm long constriction for SK-BR-3 and MDA-MB-231 cells, respectively. The cell path trajectories through the micro channels with constrictions were macroscopically scrutinized and similar behavioral patterns were observed for both cell lines and for two different pressure set points, as illustrated in Figure 2a for SK-BR-3 cells. A first observation is the large dispersion (i.e. over two orders of magnitude) of the “total transit time”, i.e. the total time spent in the constricted channels, whatever the applied pressure set point (Figure 2b). Quite interestingly, these “position versus time graph” curves revealed that the main factor limiting migration is the arrest in the first constriction. Once this constriction is passed, the subsequent ones are crossed smoothly with minimal arrest (Figure 2b). Both cell types undergo a strong deformation in the first constrictions (Figure 2d), which increases with the constriction length as expected from a crude volume conservation hypothesis. This initial deformation is partly maintained in the rest of the cell journey through the subsequent constrictions (Figure 2a). Quite interestingly too, the cell residence time in the first constriction differs between cell types, with mean values of 116 ms for SK-BR-3 cells and 34 ms for MDA-MB-231 cells (Supplementary Table 1). The ratio between the cell residence time in the first constriction and the total transit time is plotted in Figure 2c, showing that around 50% of the total transit time is spent in the first constrictions in all constricted geometries. Let us note that, however, only MDA-MB-231 cells display a significant increase of residence time as a function of the length of the constriction. Overall, these results suggest that cancer cells crossing constrictions retain a persistent memory of such an event, which also seems to depend on the initial cell type. We then explored the consequences of this morphological plasticity observed at the cellular, subcellular and molecular levels. We focused our efforts on the nucleus, the biggest organelle in the cell, which was previously identified as the limiting one for cells in migration through narrow constrictions.^23^

**Figure 1.**
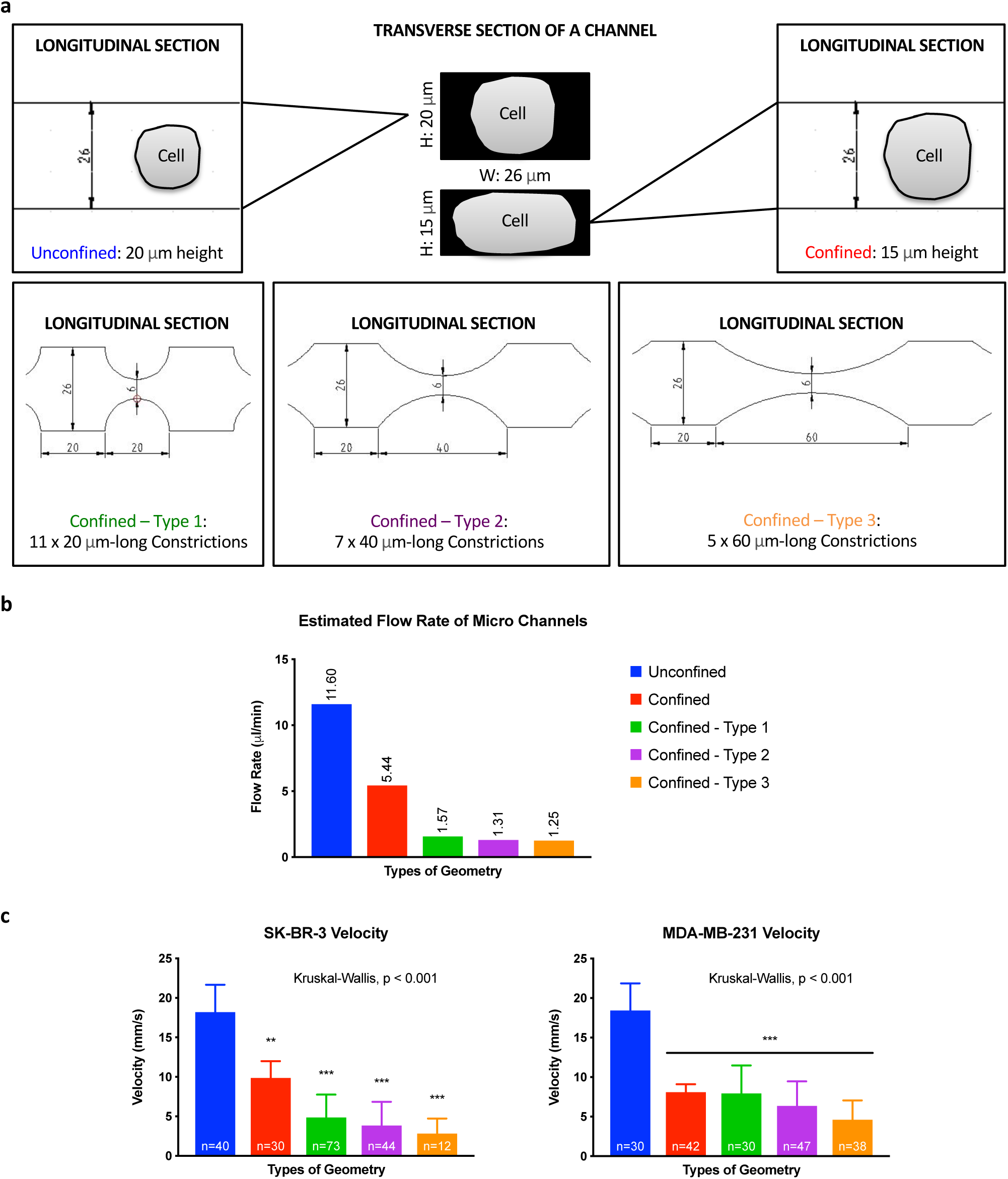
Hydrodynamics and velocity measurements in the microfluidic system at a constant applied pressure of 10 kPa. **(a)** Schematic designs of microfluidic channels without and with micro constrictions. **(b)** Estimated flow rates of various microfluidic channels. **(c)** Velocities of SK-BR-3 (left) and MDA-MB-231 (right) cells transiting through various microfluidic channels. Error bars represent standard deviation. Dunn’s multiple comparisons post hoc test. *, significance level at 0.033; **, significance level at 0.002; ***, significance level < 0.001; ns, not significant.

**Figure 2.**
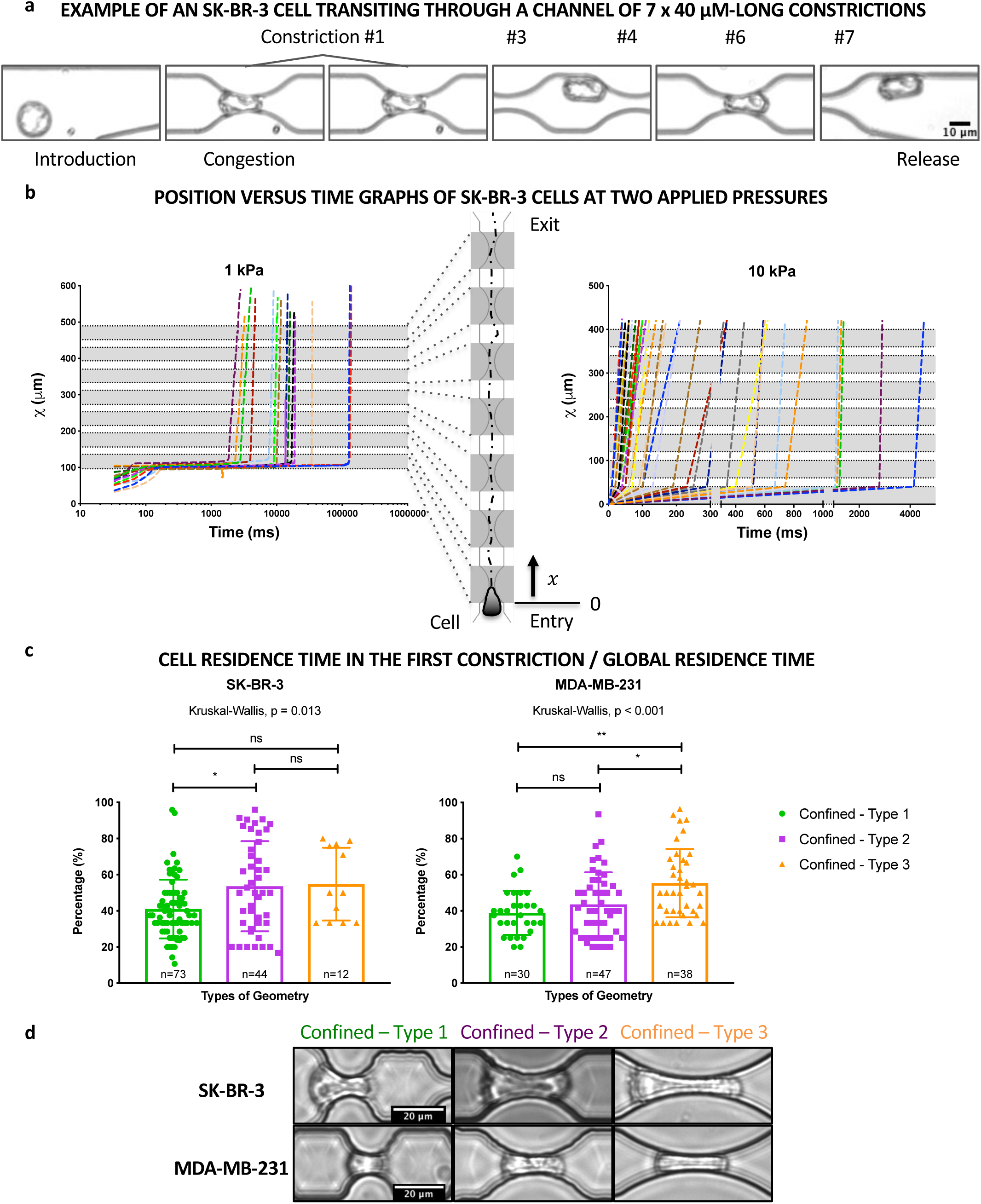
Transit and residence times of metastatic breast cancer cells in the microfluidic system at a constant applied pressure of 10 kPa. **(a)** Example of an SK-BR-3 cell transiting through a microfluidic channel with constrictions. **(b)** Position versus time graphs of SK-BR-3 cells transiting through the same type of microfluidic channel at 1 (left) and 10 (right) kPa. **(c)** Percentages of cell residence time in the first constriction for SK-BR-3 (left) and MDA-MB-231 (right) cells in three types of microfluidic channels with micro constrictions. Error bars represent standard deviation. Dunn’s multiple comparisons post hoc test. *, significance level at 0.033; **, significance level at 0.002; ***, significance level < 0.001; ns, not significant. **(d)** Brightfield images of SK-BR-3 and MDA-MB-231 cells residing in the three types of micro constrictions.

### Effects of Circulation, Confinement and Constrictions on the Morphology of Metastatic Breast Cancer Cells and Their Genome Integrity

First, we characterized the nuclear morphological features of single tumor cells after being subjected to confinement and constrictions under flow. Briefly, SK-BR-3 and MDA-MB-231 cells were isolated from the microfluidic system, fixed while in suspension, and their nuclei stained with DAPI (Figure 3a). Approximately, one thousand cells were collected in one hour. Therefore, the maximum time over which cells are left in suspension, including the time spent before and after crossing the constrictions, is under an hour. Since each of the five geometric microfluidic models could impose unique mechanical constraint on the cells, their cell area, nucleus area, nuclear to cytoplasmic (N:C) ratio, and nucleus aspect ratio (AR) were quantified for each geometry (Figure 3b). Morphological analysis showed significant increases of nucleus areas in SK-BR-3 cells in most conditions, with the highest changes observed as a result of circulation without confinement (channel height of 20 μm) and with confinement (channel height of 15 μm). This suggests that these cells respond strongly to being suspended and submitted to shear stress, and relatively less to the additional strain provided by constrictions. Conversely, MDA-MB-231 cells appeared quite insensitive to the transition from an adherent to a suspended state, and from being circulated. They however display significant changes in nucleus size when constricted. These trends were similar regarding cell size. This led to almost unchanged nuclear to cytoplasmic size ratio for both cell types and conditions, except for the unconfined and confined conditions for SK-BR-3 cells reflecting the large observed relative changes of nucleus area in this cell type. Finally, the nucleus aspect ratio (computed from an ellipsoidal fit, see Methods) appeared significantly higher in MDA-MB-231 cells, as already reported by Xia et al.^16^, but was not affected by any of the stress encountered by the cells. Subsequently, to further investigate how the morphological changes observed in most circulated conditions as compared to the adherent control condition affect the genome integrity of the cells, the extents of DNA damage and repair activity due to circulation, confinement and constrictions were investigated by staining the cells with γ-H2AX (Phospho-Ser139) antibody, a marker for DNA damage response (Figure 4a). The integrated densities of γ-H2AX in the nucleus and cytoplasm fractions were measured and their ratio were represented (N:C ratio, see Figure 4b). These results showed that DNA damage and repair activity is significantly activated in SK-BR-3 cells in all circulated groups, as compared to the uncirculated (control) group. In MDA-MB-231 cells, DNA damage and repair activity is constitutively activated (N:C ratio > 1) even in the control, and appears to be increased with a statistical significance only in the “unconfined” circulated case. Further analysis will be needed to understand this point but we can note that a similar trend (also not statistically significant) is observed for the N:C ratio in size.

**Figure 3.**
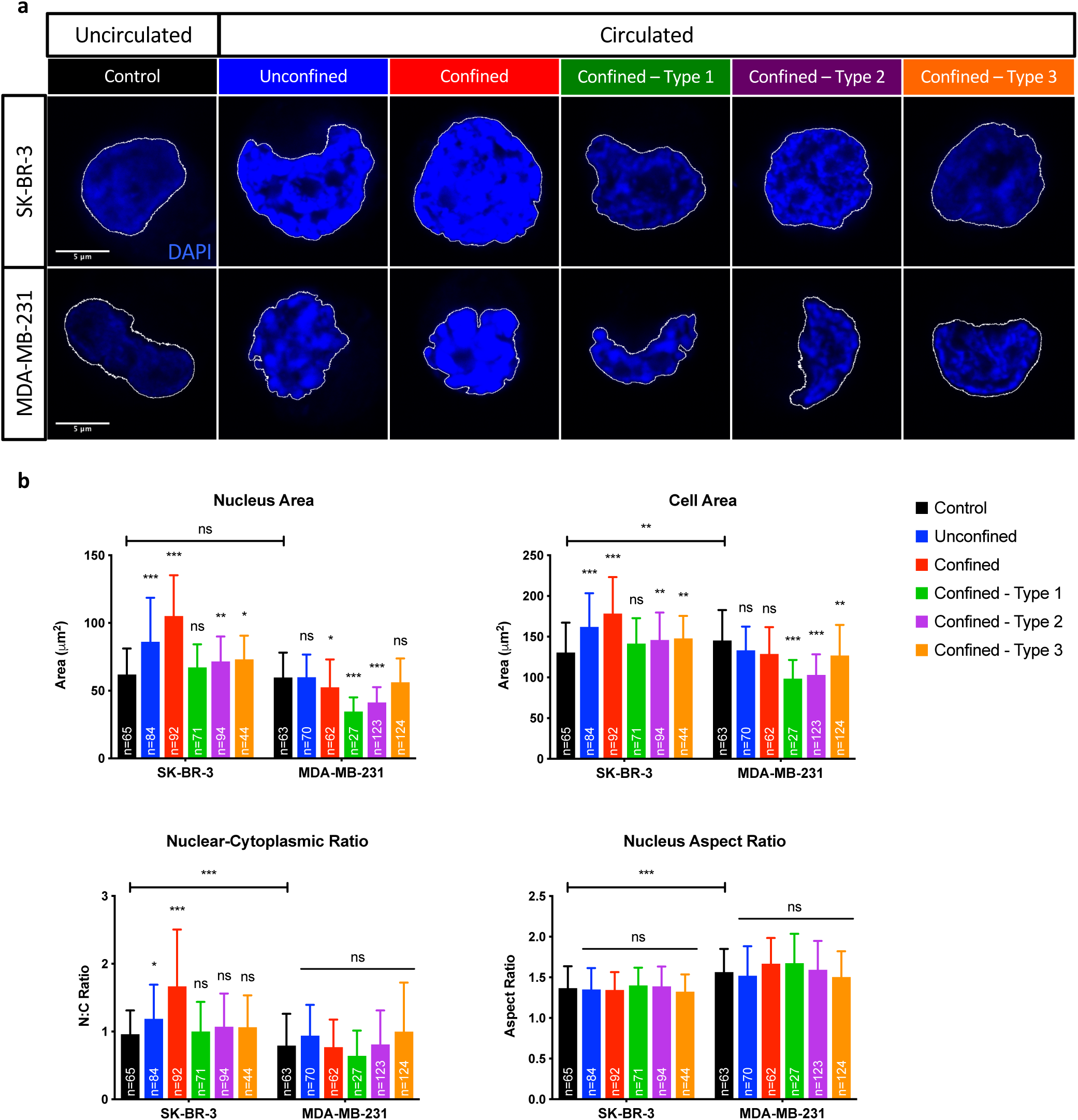
Morphological analysis of SK-BR-3 and MDA-MB-231 cells post-circulation, -confinement and –constrictions at a constant applied pressure of 10 kPa. **(a)** DAPI staining of SK-BR-3 (top row) and MDA-MB-231 (bottom row) cells. **(b)** Quantifications of nucleus area, cell area, N:C ratio and nucleus AR of SK-BR-3 and MDA-MB-231 cells. Error bars represent standard deviation. Mann-Whitney U test, two-tailed used to compare control groups between cell types. Dunn’s multiple comparisons post hoc test. *, significance level at 0.033; **, significance level at 0.002; ***, significance level < 0.001; ns, not significant.

**Figure 4.**
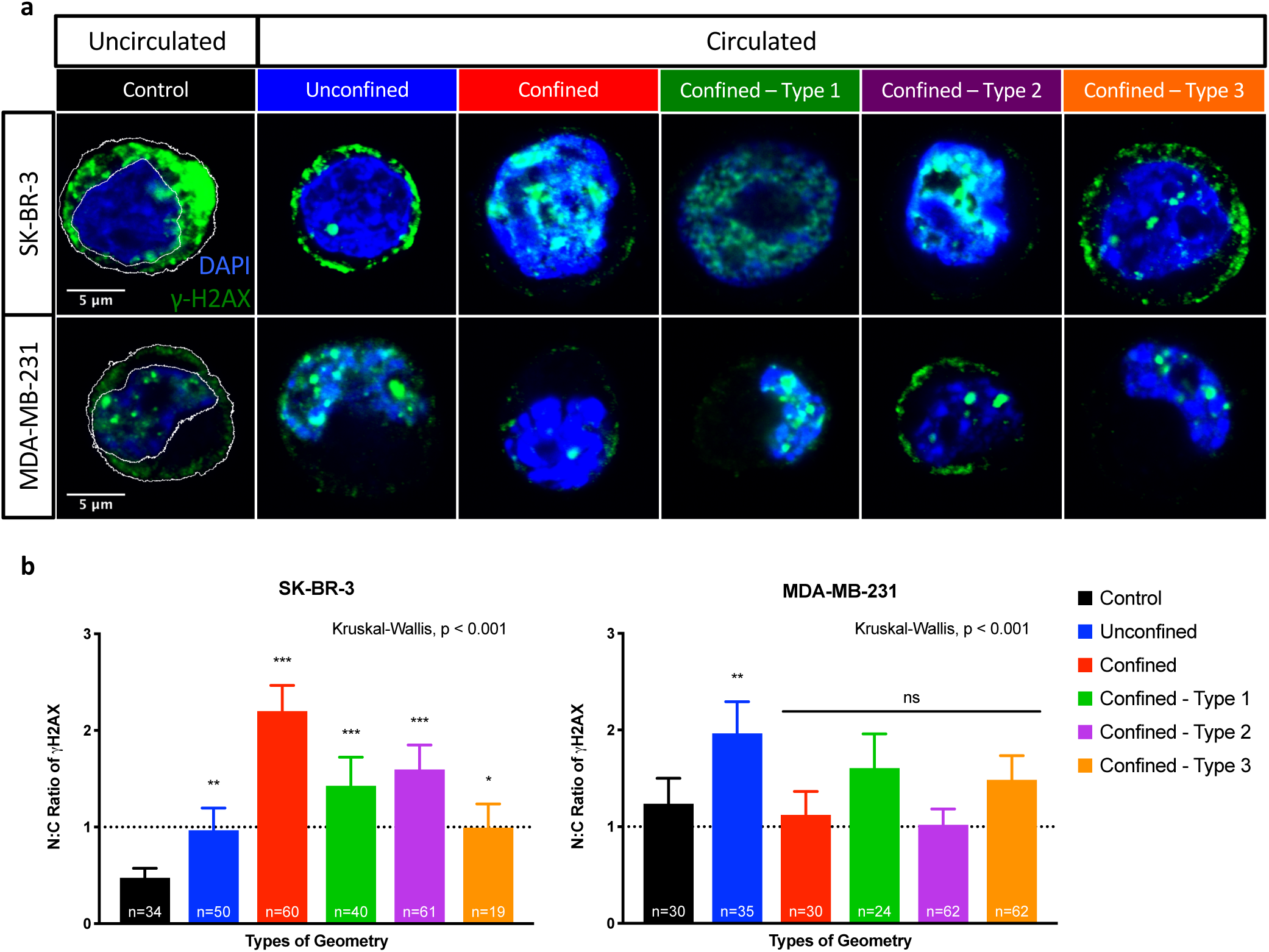
Genome integrity analysis of SK-BR-3 and MDA-MB-231 cells post-circulation, -confinement and -constrictions at a constant applied pressure of 10 kPa. **(a)** DAPI (blue) and γ-H2AX (green) staining of SK-BR-3 (top row) and MDA-MB-231 (bottom row) cells. **(b)** Quantifications of nuclear to cytoplasmic ratios of γ-H2AX integrated densities. Cytoplasmic co-localization – N:C ratio < 1; nuclear co-localization – N:C ratio > 1. Error bars represent standard deviation. Dunn’s multiple comparisons post hoc test. *, significance level at 0.033; **, significance level at 0.002; ***, significance level < 0.001; ns, not significant.

### Gene Expression Analysis of Key Regulators of Epithelial-to-Mesenchymal Transition

Next, a panel of EMT transcriptional signatures were investigated for possible induced changes in tumor cells due to circulation, confinement and constrictions. Quite interestingly, significant changes in gene expression of EMT transcriptional signatures were observed in response to circulation alone (Figure 5). These changes are associated with an increase of expression in SK-BR-3 cells, 5 over 9 genes being significantly up-regulated (E-cadherin, Snail1, Twist2, ZEB1 and ZEB2). For MDA-MB-231 cells, a similar yet less pronounced trend was observed (i.e. only Snail1 and ZEB2 were significantly overexpressed), and notably, down-regulations of N-cadherin, Twist1 and Twist2 were observed. When the effects of confinement and constrictions were added on top of circulation, no significant changes were detected for most of the transcriptional signatures investigated for both cell lines. SK-BR-3 cells however, displayed significant down-regulated genes for Vimentin and Snail2 when compared to the unconfined circulated condition (Figure 6a). Quite a different situation was observed for MDA-MB-231 cells. Amongst the nine EMT transcriptional signatures tested, only E-cadherin was significantly down-regulated while Twist2 revealed a striking increase in relative expression for all conditions (Figure 6b). Since Twist2 is known to play a role in tumor progression in breast cancer^24^, even though its expression has been less explored than that of Twist1, we decided to measure its expression and cell localization.

**Figure 5.**
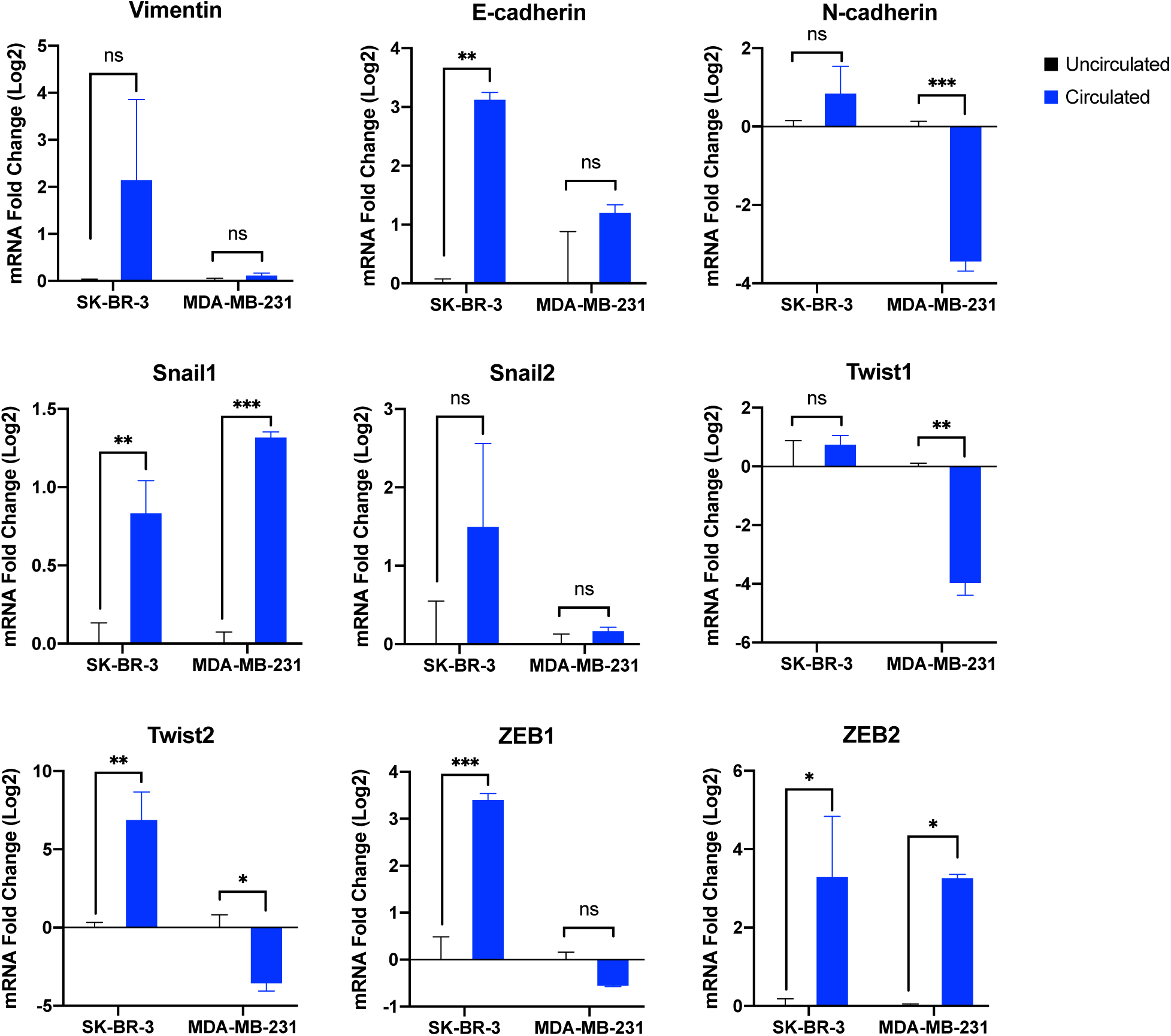
Relative expression (log2, mean ± s.e.m.) of Vimentin, E-cadherin, N-cadherin, Snail1, Snail2, Twist1, Twist2, ZEB1 and ZEB2 tested in two metastatic breast cancer cell types (n=3 per transcript). Holm-Sidak multiple comparisons post hoc test used to compare relative expression of all transcripts in cells in the uncirculated group with the circulated group. *, significance level at 0.033; **, significance level at 0.002; ***, significance level < 0.001; ns, not significant.

**Figure 6.**
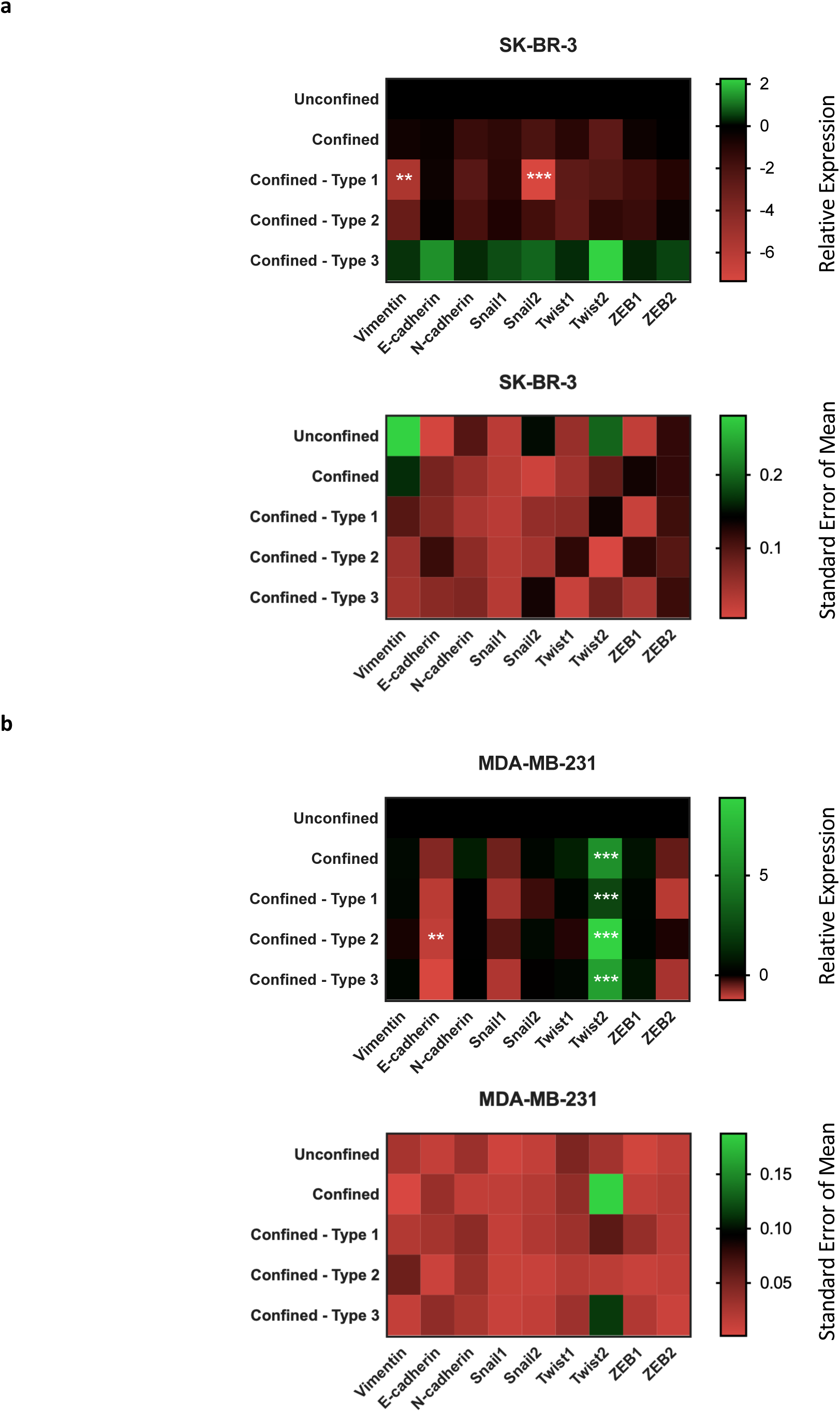
Heat map illustration of relative expression (log2, mean) and (standard error of mean) of Vimentin, E-cadherin, N-cadherin, Snail1, Snail2, Twist1, Twist2, ZEB1 and ZEB2 tested in two metastatic breast cancer cell types (n=3 per transcript). Holm-Sidak multiple comparisons post hoc test used to compare relative expression of all transcripts in cells in the confined groups with the unconfined group. *, significance level at 0.033; **, significance level at 0.002; ***, significance level < 0.001; ns, not significant. **(a)** SK-BR-3 cells. **(b)** MDA-MB-231 cells.

### Co-localization of Twist2 Protein

The co-localization of Twist2 protein in the nucleus and cytoplasm was determined in SK-BR-3 and MDA-MB-231 cells in response to fluid shear stress alone, and to the combination of fluid shear stress and mechanical constraints by staining with Twist2 antibody (Figure 7a). The integrated densities of Twist2 in the nucleus and cytoplasm fractions were measured and represented as N:C ratio (Figure 7b). SK-BR-3 cells in the circulated groups exhibited significant increases of nuclear co-localization of Twist2 as compared to the control group. On the other hand, MDA-MB-231 cells in the circulated groups exhibited either maintained N:C levels of Twist2 localization or significant increases of cytoplasmic co-localization of Twist2 in two of the constricted conditions as compared to the control group.

**Figure 7.**
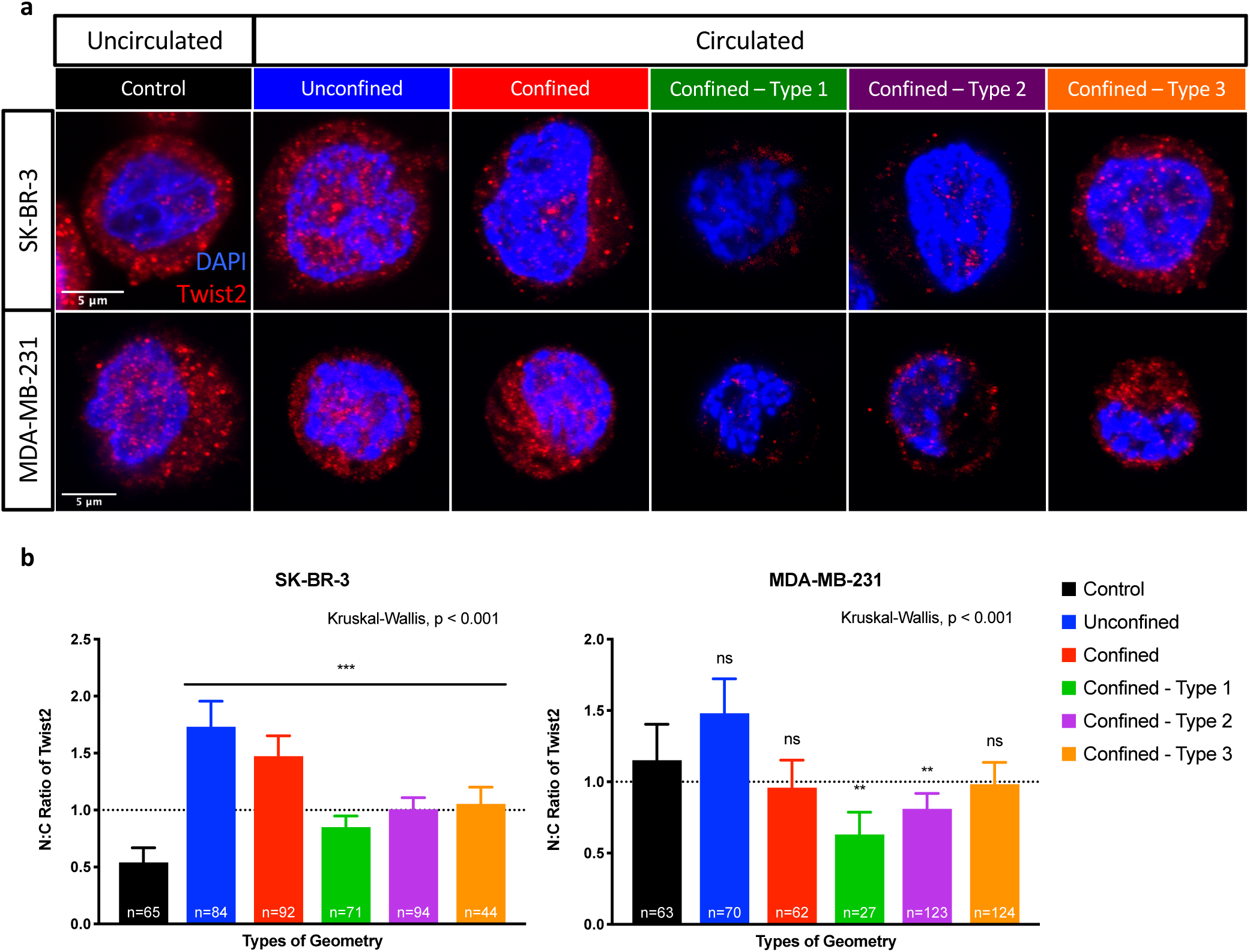
Immunofluorescent analysis of SK-BR-3 and MDA-MB-231 cells post-circulation, -confinement and -constrictions at a constant applied pressure of 10 kPa using Twist2 antibody. **(a)** DAPI (blue) and Twist2 (red) staining of SK-BR-3 (top row) and MDA-MB-231 (bottom row) cells. **(b)** Quantifications of nuclear to cytoplasmic ratios of Twist2 integrated densities. Cytoplasmic co-localization – N:C ratio < 1; nuclear co-localization – N:C ratio > 1. Error bars represent standard deviation. Dunn’s multiple comparisons post hoc test. *, significance level at 0.033; **, significance level at 0.002; ***, significance level < 0.001; ns, not significant.

## Discussion

Circulation of detached cancer cells within PDMS microchannels provides a foremost model to comprehend the fate of CTCs during their transient existence into the blood circulation. In the present work, we have studied several aspects of this fate, starting from the dynamical behavior of cells that were flow-driven into constrained microchannels. Our data highlight differential mechanical characteristics of poorly versus highly metastatic cancer cell lines in terms of deformability and plasticity. In the literature, the mechanical phenotype of cells is emerging as a potential biomarker for cell types, particularly cancer cells. Many studies have reported the usage of microfluidic devices to measure the mechanical phenotypes of cells including transit time, entry time, cell size, elastic modulus and cell fluidity.^12, 25–27^ This was applied to dissociated cells from tumors, in line with previously proposed deformation assays.^28, 29^ We believe that this is particularly interesting for CTCs, since they are expected to undergo large mechanical constraints during their crossing of microcapillaries *in vivo*. In the present study, not surprisingly, higher confinements led to lower average velocities for cells transiting across a channel with micro constrictions (Figure 1). Cells undergoing flow-induced migration under 3D confinement with multiple constrictions generally underwent a significant arrest at the first constriction (Figure 2). Subsequently, they used up a significant amount of time to cross this first encounter. We interpret these high residence times in the first constriction as the time needed by the cells to deform in order to conform to the mechanical constraint of the constriction and to cross it. Once deformed by this first constriction, their subsequent crossings of constrictions of the same type were significantly faster. This suggests a capacity to store mechanical deformation on timescales comparable to the transit time between constrictions.

While both cell types studied here are of epithelial breast cancer origins, they exhibit morphological differences in 2D culture: SK-BR-3 cells display grape-like morphologies and MDA-MB-231 cells display stellate morphologies. This difference was reflected pronouncedly in the nuclear morphological analysis between the cell types post-circulation (Figure 3). Prior to flow, both cell types SK-BR-3 and MDA-MB-231 have similar nucleus areas (P=0.520). The nucleus is the densest organelle of a cell, and its mechanical resistance should be a strong hindrance to the crossing of constrictions. Therefore, one may expect that both cell types have similar responses in terms of crossing time. Interestingly however, both cell types responded inversely to confinement regarding changes in the projected cellular areas: SK-BR-3 displays significant increases in nucleus and cell areas, while MDA-MB-231 displays significant opposites. This could explain the higher values of transit times with SK-BR-3 cells, as compared to MDA-MB-231 cells (Supplementary Table 1). The higher average transit times with SK-BR-3 cells as compared to MDA-MB-231 cells could also be explained by significant differences in initial nucleus aspect ratio between both cell types (mean nucleus AR of 1.57 and 1.37 for MDA-MB-231 and SK-BR-3 cells, respectively, P<0.001), as already proposed by Xia et al. in a similar context (i.e. the comparison between MDA-MB-231 and MCF-7 cells, the latter displaying a relatively lower aspect ratio).^16^ At similar volume, a rounder morphology requires a cell to undergo more deformation to successfully cross constrictions as compared to a more elongated morphology. Fascinatingly, too, the variations in nuclei and cell morphological parameters between cell types (Figure 3) are significant yet relatively modest but their consequences on the transit time for the first constriction are more dramatic, with mean values higher by a factor 3 to 7 in SK-BR-3 as compared to MDA-MB-231 cells (Supplementary Table 1). This might suggest that the mechanical response of the cell, considered as a viscoelastic body, is strongly nonlinear, and display “hardening” (higher apparent modulus) when deformation increases. This raises intriguing and, to our knowledge, not fully understood questions regarding the mechanics of the nucleus.^25, 27^ Different experimental tools would be needed to address this question in detail, as it is difficult to characterize different cell samples based on transit time data only.^30^

Studies on DNA damage response (DDR) to mechanical stress, particularly fluid shear stress, mechanical confinement and constrictions are still scarce,^31^ yet they are slowly emerging within the last years. DDR is a series of coordinated responses designated to remove damage incurred to the genome.^32^ Using transformed cell lines, Singh et al. reported that lamins A/C, essential nuclear envelope proteins, are required for maintaining genomic stability and that their depletion stalls DDR.^33^ Later, Davidson et al. associated the role of lamins A/C and DNA damage with migration-induced nuclear deformations in fibroblasts.^23^ Several additional studies emerged, reporting that the genomic instability caused by migration-induced nuclear deformation and DNA damage promotes cancer heterogeneity.^34–36^ Furthermore, Jacobson et al. reported and identified changes in transcripts from RNA-sequencing of neutrophil-like cells in response to migration and constriction stress.^37^ This establishes an interesting interplay between nuclear mechanics, genome integrity and phenotypic transformation. Here, tumor cells that underwent circulation, confinement and constrictions indeed were found to present increased levels of DNA damage marker, γ-H2AX (Figure 4). As compared to MDA-MB-231 cells, SK-BR-3 cells exhibited more significant increases in nuclear co-localization of γ-H2AX. Simplistically, this could suggest that DDR is more weakly activated by physical mechanical constraints in MDA-MB-231 than in SK-BR-3 cells. Upon closer inspection, however, DDR is more than twice higher in MDA-MB-231 (mean N:C ratio: 0.474) than in SK-BR-3 (mean N:C ratio: 1.24) cells at baseline, i.e. prior to flow. COSMIC, the Catalogue of Somatic Mutations in Cancer (http://cancer.sanger.ac.uk) reported 4 gene mutations in SK-BR-3 cells and a whopping 174 gene mutations in MDA-MB-231 cells.^38^ This may imply that DDR mechanisms are more active in MDA-MB-231 than in SK-BR-3 cells due to a higher number of gene mutations, already in the absence of stress. The specific contribution of cell deformation to DDR thus may appear smaller in relative value in MDA-MD-231, simply because other non-mechanical causes of activation are more active. Our results, however, may also reflect a more fundamental property of mesenchymal-like cells, i.e. a better ability to deform without inducing further DNA damage.

The plasticity of EMT in cells from aggressive tumors enables them to switch from proliferative to invasive phenotypes and vice versa.^39, 40^ It has been vastly observed in mouse mammary tumor models that the initiation of EMT can occur during the early stages of tumorigenesis and progress during its later stages.^41–44^ The EMT process is considered as a hallmark-facilitating program as well as having influence on pathways linked to tumor progression and metastatic dissemination. This raises the question of the status of CTCs amongst the spectrum of functional and morphological characteristics, and notably the epithelial and mesenchymal ones. It was proposed that CTCs have had undergone EMT or are still continuously undergoing the transition while in the circulation.^45^ While it has been reported at the cellular level that the functional roles of the cytoskeleton modulate suspended cell mechanics and that of substrate-adhered cells,^11^ here we would like to address how transformations from mechanical stress, particularly that of fluid shear stress and mechanical constraints undergone by CTCs in the circulation, may induce changes in cells at the molecular level. Imani et al. recently carried out a meta-analysis on EMT transcription factors (EMT-TFs) studies reported on metastatic breast cancer and highlighted Twist1, Snail1, Snail2, and ZEB1 as key EMT-TFs that are involved in metastatic breast cancer through different signaling cascades.^46^ Here, we showed that, in SK-BR-3 and MDA-MB-231 cells, there is a substantial relationship between EMT-TFs and mechanical stress from fluid shear stress and mechanical constraints mimicking those present in the microcirculation – the route that CTCs take in order to disseminate (Figure 5). Amongst the EMT markers tested, the ZEB family, Twist1 and N-cadherin did not show any significant difference, while Vimentin, Snail2 and E-cadherin genes were significantly down-regulated. Further work is necessary for a fully consistent explanation of these first results. Some significant pictures arise, however, notably, for MDA-MB-231 cells, a systematic overexpression in all confined conditions of Twist2, a nuclear protein playing the role of distinct tissue-restricted transcription factor (Figure 6).^47^ The central role of Twist2 in embryogenesis and mesodermal development, and its targeting of multiple genes coding for cell-fate proteins inevitably links them to cancer and oncogenesis.^48^ This gene has also been reported to be involved in metastasis formation through EMT, thereby facilitating cancer cell invasion in epithelium-based cancers.^49, 50^ It was also reported that the EMT program might be activated transiently through nuclear Twist2 in the tumor invasion front to facilitate cancer cell invasion and metastasis.^24^ An overexpression of Twist2 in circulated MDA-MB-231 shed light on the singularity of Twist2 as compared to Twist1, and its possible activation under certain conditions of stress. In our case, this refers to shear stress. In addition to this gene expression pattern, we observed heterogeneous co-localizations of Twist2 protein between the nucleus and the cytoplasm (Figure 7). At baseline, Twist2 was co-localized much more in the cytoplasm of SK-BR-3 (mean N:C ratio: 0.541) than that of MDA-MB-231 (mean N:C ratio: 1.151) cells. While significant decreases of N:C ratio of Twist2 in MDA-MB-231 cells flowing into constrictions could be put in connection with the significant decrease in nucleus and cell areas (Figure 3), significant increases of N:C ratio of Twist2 in SK-BR-3 cells for all circulation experiments (P<0.001) highly suggest a link between nuclear Twist2 and EMT in circulating SK-BR-3.

In this study, it was observed that epithelial-like SK-BR-3 cells underwent more stark changes in (1) cellular behavior, (2) morphological features and (3) DDR mechanisms, as compared to mesenchymal-like MDA-MB-231 cells after exposure to mechanical stress from fluid shear stress and mechanical constraints. The epithelial versus mesenchymal status of these cells might contribute to the difference in response to applied mechanical stimuli between cell types. SK-BR-3 cells exhibit epithelial-like morphologies (which suggests higher rigidity) while MDA-MB-231 cells exhibit mesenchymal-like morphologies, which suggests higher deformability. These findings suggest that circulation may have a selective effect on a population of CTCs with a pre-existing heterogeneity, e.g. in terms of survival (more detrimental to epithelial-like phenotype), penetration in organs capillaries (favoring the more flexible mesenchymal phenotype), and/or cell arrests leading to extravasation (oppositely favoring the more rigid epithelial one).

In conclusion, flowing cancer cells from cell lines in micro channels, in conditions tailored to mimic those encountered in the blood flow, has shown significant impacts on cellular processes that can lead to changes in morphological features and genetic expressions. More specifically, this circulation process encompasses (1) the hydrodynamic shear stress from blood flow and (2) the mechanical deformations from the geometrical constrictions of the microvasculature. It was shown that changes in morphological features and genetic expressions from this circulation process differ from one cell type to another. Notably, two cell lines presenting an epithelial-like and mesenchymal-like state, respectively, have different responses to different mechanical stress. Therefore, this work strongly suggests that the mechanical process of circulation in the blood flow may have a significant effect on cells at morphological and gene expression level, and that this effect is cell type-dependent.

Since circulation in the blood is a necessary step in a large fraction of metastatic dissemination events (except for dissemination through the lymphatic system), these findings suggest that this step should not be ignored when trying to comprehend the metastatic process as a whole. Future research efforts should involve (1) an increase in the number of constriction encounter events, through the courtesy of a more elaborate microfluidic design, (2) a more detailed study of the morphological recovery of cells post-constrictions, (3) potential DNA damage repair processes, (4) a comparison of the migratory potential of cells before and after transformation by the circulation and constrictions, and (5) the use of CTCs from patients. This latter study is also challenging, due to the rarity of CTCs. Of course, these studies should be performed in light of and in parallel with the current evolution of the general understanding of the instability of cancer cells and CTCs, and notably the EMT and MET. This should help to achieve a more in-depth understanding of the molecular mechanisms activated or repressed in CTCs in the blood microcirculation from hydrodynamic shear stress and mechanical deformations.

## Methods

### Photo-Masks Design

A computer-aided design software application for two-dimensional design and drafting called QCAD was used to design different types of geometry (Supplementary Figure 1). The total length of one geometrical channel is 420 µm. Each type of design consists of four 420 µm-long channels that are arranged in parallel. A control design without constrictions has a channel width of 26 µm. All constrictions have a width of 6 µm at its narrowest. three types of designs with different constrictions lengths were prepared. The first type contains 11 constrictions with each measuring 20 µm in length. The second type contains 7 constrictions with each measuring 40 µm in length. The third type contains 5 constrictions with each measuring 60 µm in length. The constrictions are connected with a transit channel 20 µm by 26 µm in dimension.

### Optical Photolithography

The above two-dimensional geometrical designs were etched onto a glass substrate with a chrome layer. The production of this chrome mask was outsourced from Selba S.A., Switzerland. From this chrome mask, a silicon master mold was fabricated in a cleanroom where provisions were made to reduce particulate contamination and control other environmental parameters such as temperature, humidity and pressure (Supplementary Figure 2). A silicon wafer was used as a substrate. A negative photoresist SU-8 is spin-coated onto the substrate. SU-8 2010 photoresist was used to obtain structures with structure heights of 20 µm and 15 µm. After spin-coating the substrate with SU-8 2010 photoresist, the photoresist-coated substrate is soft baked at 95°C (Supplementary Table 2a). An optical lithography mask aligner called MJB4 (SÜSS MicroTec AG, Germany) was used to expose UV light unto the chrome mask that was aligned on the photoresist-coated substrate. The UV light exposure dosage was optimized for each of the desired thickness (Supplementary Table 2b). Post exposure bake at 95°C is carried out directly after exposure. The exposed photoresist was then developed in a shaking bath of photoresist developer (Supplementary Table 2c). After development, the developed image was rinsed with fresh developer solution for approximately 10 seconds, followed by a rinse with Isopropyl Alcohol (IPA) for another 10 seconds. Then, it was air-dried with pressurized air and hard baked for 30 minutes at 300°C. The fabrication of a silicon master mold is now complete.

### Soft Lithography

From this silicon master mold, a microfluidic polydimethylsiloxane (PDMS) chip is fabricated through soft lithography (Supplementary Figure 3). PDMS silicon elastomer, Sylgard 184 was purchased from Dow Corning (Michigan, USA). It was mixed with the provided curing agent at a ratio of 9:1 and then poured over the silicon master mold. They were cured at 70°C and left to harden for 4 hours. After curing, the PDMS stamp was separated from the silicon master mold. 1.5 mm holes were punctured on the designated areas of the PDMS stamp in order to create reservoirs. Oxygen plasma treatment was performed to induce permanent bonding and the chip was closed by a glass coverslip. From this, PDMS-bonded slides were used as chips for the microfluidic system.

### Microfluidic System

All reagents and cell suspensions used during the experiments (Supplementary Figure 4a) were stored in pressurized containers that were connected to the microfluidic chip through a manifold valve (Fluigent, France). An MFCS-8C Flow Controller (Fluigent, France) was used to control the pressure independently in each container to drive all reagents and cell suspensions into the microfluidic chip. A heat generator was used to warm the microfluidic device at 37°C throughout the experimental duration in order to achieve optimal culture conditions for experiments with live cells. Each microfluidic chip experiment experienced three major stages (Supplementary Figure 4b). The first stage is the priming of the chip in which the microfluidic channels were sterilized with 70% Ethanol (EtOH) followed by a washing step with 1X Phosphate-Buffered Saline (PBS). In order to have non-adhesive surfaces, the channels were coated with 100% of 1 mg of PLL-g-PEG (Susos AG, Switzerland) solution that was dissolved in 100 mM of sodium bicarbonate (NaHCO3) buffer at pH 8.5 for 1 hour.1 The channels were washed with 1X PBS and then filled with serum-free Dulbecco’s modified Eagle’s medium (DMEM) with HEPES (ThermoFisher Scientific, USA). The intermediate stage is the introduction of cells into the chip. A suspension volume of approximately 5000 cells in serum-free DMEM with HEPES was introduced into the microfluidic chip. This approximation of cell number was deemed optimal for the microfluidic models used in order to optimally observe single cell flow-induced migration in the channels without cell clumping or channel clogging. Here, single cell flow-induced migration was recorded based on the image acquisition control of frame rate. The image acquisitions of cells crossing the micro channels of constrictions 50 frames per second were produced using the Basler acA800-510um USB 3.0 camera (Basler AG, Germany). The final stage is the retrieval of cells from the chip. Following the introduction of cells into the microfluidic chip, circulation of cells was allowed for 1 hour. Then, the cells were carefully retrieved from the output reservoir for two follow-up tests. The final volume of retrieved cells from one microfluidic experiment was approximately 10 µL. The first follow-up test was for molecular genetic analysis and the second follow-up test was for immunofluorescence (IF) analysis of cellular proteins.

### Cell Tissue Culture

All cell lines were purchased from American Type Culture Collection (Virginia, USA). All cell culture reagents were purchased from Gibco, ThermoFisher Scientific (Massachusetts, USA). Human breast cancer cell lines: SK-BR-3 and MDA-MB-231; and epithelial Madin-Darby canine kidney (MDCK) II cells were cultured in complete Dulbecco’s modified Eagle’s medium (DMEM) that was supplemented with 100 U/ml aqueous penicillin, 100 mg/ml streptomycin and 10% fetal bovine serum (FBS). Cells were maintained at 37°C in a humidified atmosphere containing 5% CO2 and harvested with TrypLE (1X). Cell dissociation was deactivated with complete media. Cells were pelleted by centrifugation at 180 x g (relative centrifugal force, RCF) for 5 minutes and then re-suspended in complete media. The cell suspensions were used only when their viability as assessed by trypan blue exclusion exceeded 95% before use. Cell viability was determined and calculated as the number of viable cells divided by the total number of cells within the grids on a hemocytometer. The cell density of cell suspensions was determined using a hemocytometer. 0.1 mL of 0.4% of trypan blue solution was added to 0.1 mL of cells. The trypan blue solution and cell mixture was loaded into a hemocytometer and examined immediately under a microscope at low magnification. The number of blue staining cells and the number of total cells were counted. Cell viability should be at least 95% for healthy log-phase cultures.

### Quantitative Real-Time Polymerase Chain Reaction (qRT-PCR)

Total RNA extraction was performed using the ARCTURUS® PicoPure® RNA Isolation Kit (Applied Biosystems, USA). The kit was designed to recover high-quality total RNA consistently from fewer than ten cells and even from a single cell. Total RNA extraction was executed as directed by the manufacturer. Total RNA was measured using Qubit RNA reagents (Invitrogen, USA). Measurements were conducted using the Qubit 3.0 Fluorometer (Invitrogen, USA). All the RNA samples did not undergo more than two freeze-thaw cycles to avoid any potential nucleic acid degradation. Total RNA reverse transcription (RT) was performed using the High-Capacity cDNA Reverse Transcription Kit (Applied Biosystems, USA). After reverse transcription, the cDNAs were diluted by 10-fold before amplification. cDNA amplification was performed using KiCqStart® SYBR® Green qPCR ReadyMix™ (Merck, Germany). The mRNA primer designs (Sigma-Aldrich, USA) are listed in the Supplementary Table 3a. All real-time PCR reactions were performed in triplicates using a SmartCycler® automated real-time PCR system (Cepheid Inc., USA). Supplementary Table 3b sets the thermal cycling conditions for amplification. All target mRNA expressions were normalized to reference gene GAPDH. Relative mRNA levels were calculated using the – ΔΔCt method.

### Immunofluorescence

10 µL of cell suspensions that were retrieved from the microfluidic chip device as aforementioned were fixed with equal volume of 4% paraformaldehyde (PFA) and then dispensed on a poly-L-lysine coated glass slide (Sigma-Aldrich, USA). The cells were incubated at room temperature for 15 minutes. The fixed cells were washed with 1X PBS and then permeabilized with 0.1% Triton-X 100 at room temperature for 10 minutes. The cells were washed with 1X PBS and then blocked with 4% Bovine Serum Albumin (BSA) for 1 hour at room temperature. The cells were washed with 1X PBS and then incubated with primary antibody cocktail in 2% BSA at room temperature for 2 hours (Supplementary Table 4). The cells were washed twice with 1X PBS and then incubated with secondary antibody cocktail in 2% BSA at room temperature for 1 hour (Supplementary Table 4). The cells were washed twice with 1X PBS and then counterstained with ProLong Gold Antifade Mountant with DAPI (Invitrogen, USA). The slide was mounted with a cover slip and the cells were visualized using the Leica TCS SP8 confocal laser scanning microscopy platform (Leica Microsystems, Germany).

### Transit Time Analysis

Videos from microfluidic experiments were analyzed with ImageJ Version 1.51. The Mtrack2 plugin was installed on ImageJ. It was used to track the positions of a cell cross a channel with constrictions (Figure 2a). First, the entire image sequences were converted to mask. Then, the outline of the channel was subtracted from each image sequence. Finally, the positions of the cell were measured from each image sequence. This method automatically churned the measurements of the position vector of the cell that allowed the calculations of cell velocity and cell residence time. This method is only effective when there is only one cell crossing a channel from beginning to end. Thus, manual tracking was done on image sequences that had more than one cell crossing the channel within a single time frame.

### Cell Morphological Analysis

All acquired images were analyzed with ImageJ Version 1.51. Regions of interest (ROIs) were used to define specific parts (cell and nuclear boundaries) of an image that was processed independently. Only the pixels within any defined ROI were included in the calculations when measured. ImageJ’s set measurements function allowed the generation of the following data: area, aspect ratio and integrated density. This method was also used to measure the protein presence and its localization in a cell. Integrated Density is the product of “Area” and “Mean Gray Value”.

### Statistical Analysis

All statistical hypothesis testing was conducted using the GraphPad Prism 7 software (GraphPad Software Inc., USA). An alpha of 0.05 was used as the cut-off for significance. The Kruskal-Wallis test followed by Dunn’s multiple comparisons post hoc test was used to compare three or more independent samples of equal or different sample sizes. This method was used to compare data on velocity, transit time and residence time of cells in the microcirculation on five independent samples from the same applied pressure force. This method was also used to compare data on nucleus area, nucleus length, nuclear to cytoplasmic area ratio, nucleus aspect ratio, cell area and nuclear to cytoplasmic ratio of γH2AX and Twist2 proteins of cells post-microcirculation on six independent samples from the same applied pressure force. The two-way analysis of variance (ANOVA) followed by Holm-Sidak multiple comparisons post hoc test was used to compare relative expression of every transcript between two groups (uncirculated and circulated) of two cell types (SK-BR-3 and MDA-MB-231). This method was also used to compare relative expression of nine transcripts between two groups (unconfined and every confined) for SK-BR-3 and MDA-MB-231.

## Acknowledgements

This work was supported by the French National Research Agency (ANR) as part of the “Investissements d’Avenir” program (reference: ANR10-NANO0207; Labex and Equipex IPGG), ERCadg CellO (FP7-IDEAS-ERC-321107) and Ecole Doctorale Frontières du Vivant (FdV) – Programme Bettencourt. We are indebted to Prof. Christine Chomienne-Thomas and Dr. Benoit Ladoux for constructive comments and fruitful insights. We thank the IPGG technological platform for technical assistance and support.

## Author Contributions

H.A.C., C.V. and J.L.V. designed and conceived the study. H.A.C. planned, performed and processed the experiments, measurements and data analysis. H.A.C. wrote the manuscript. C.V. supervised and directed the project. J.L.V. contributed to supervision and interpretation of results. All authors provided critical feedback and helped shape the research, analysis and manuscript.

---

**Supplementary Figure 1.**
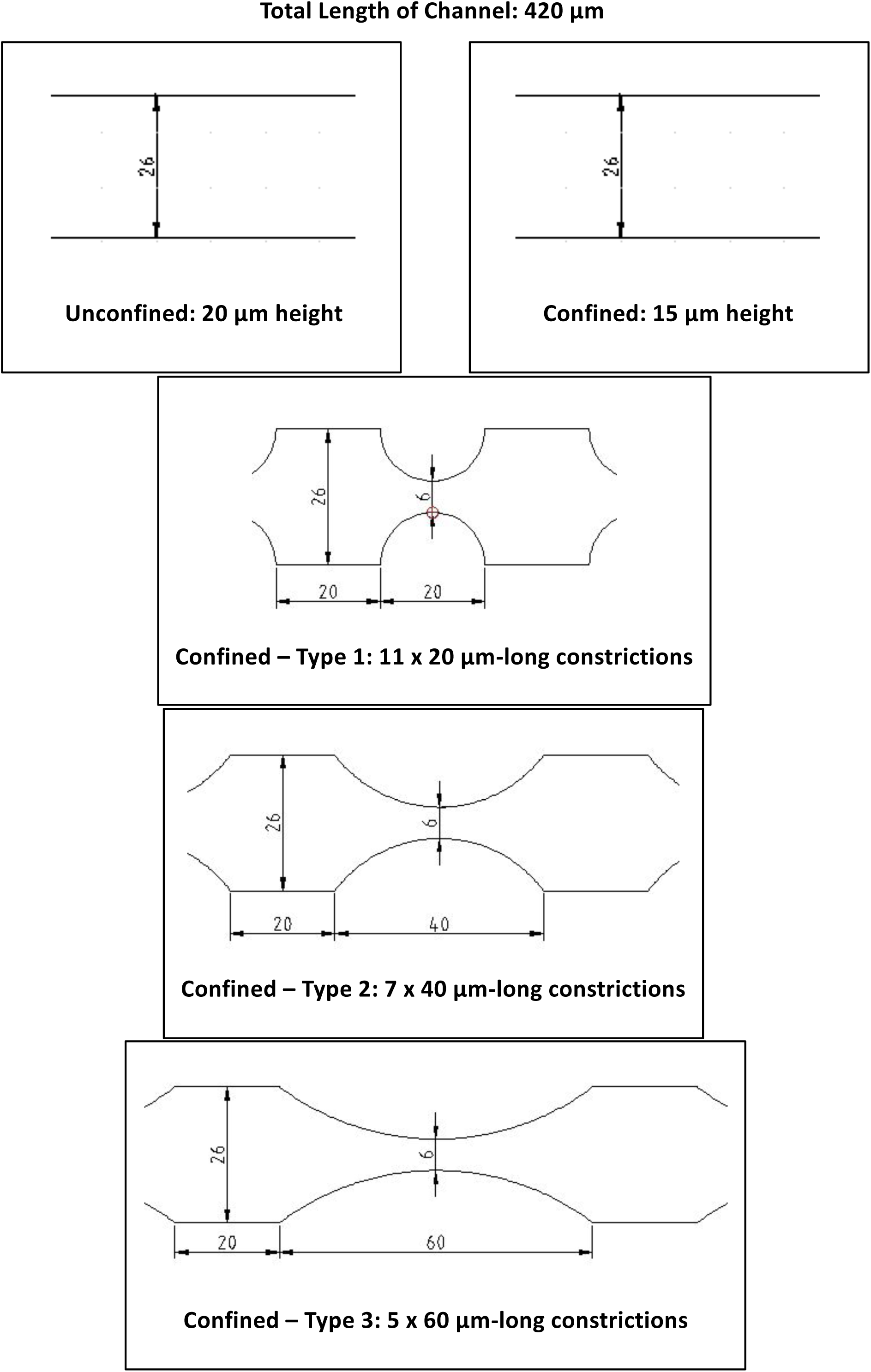
Types of two-dimensional geometrical designs. Dimensions are in micrometers.

**Supplementary Figure 2.**
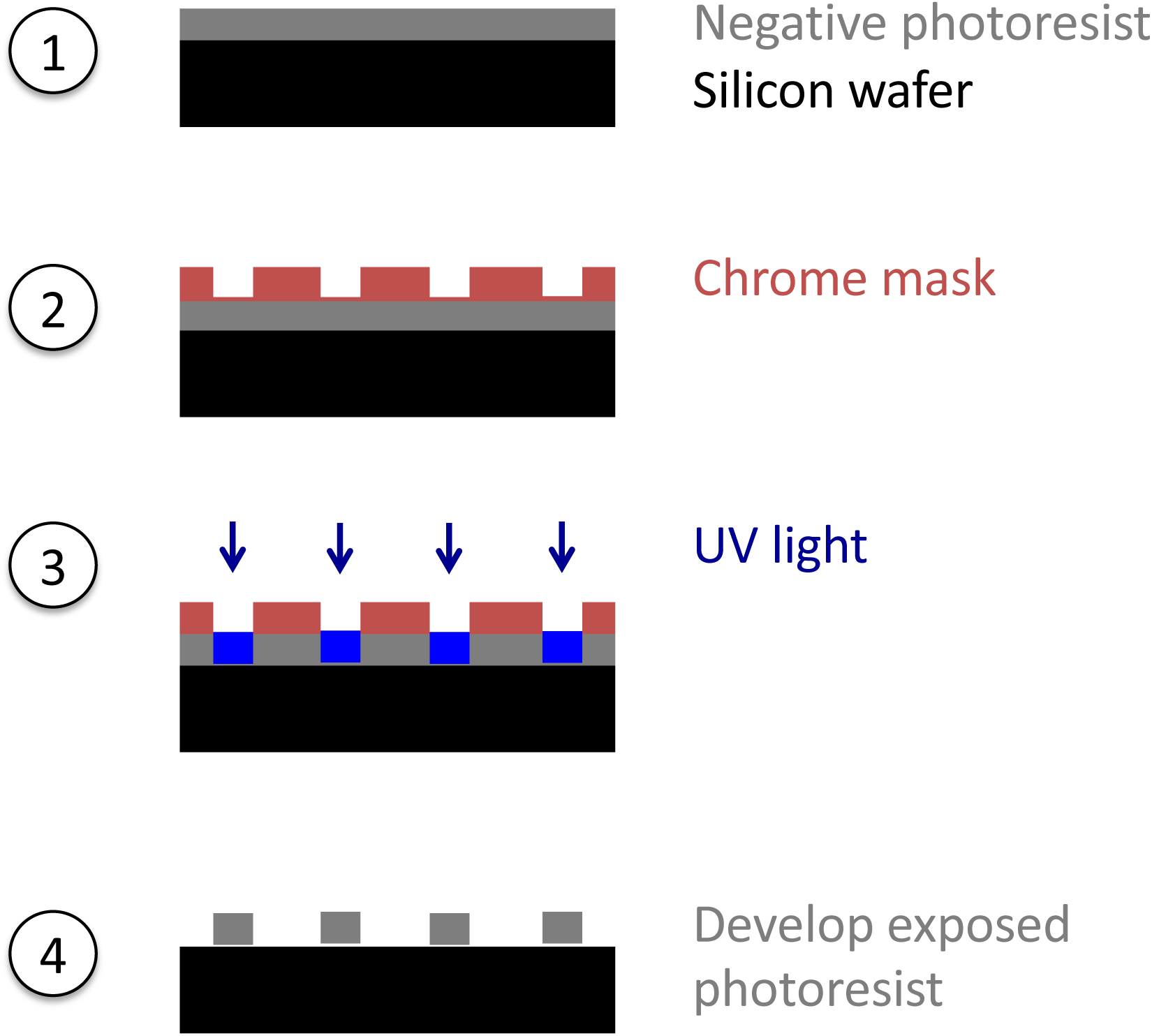
Fabrication of silicon master mold.

**Supplementary Figure 3.**
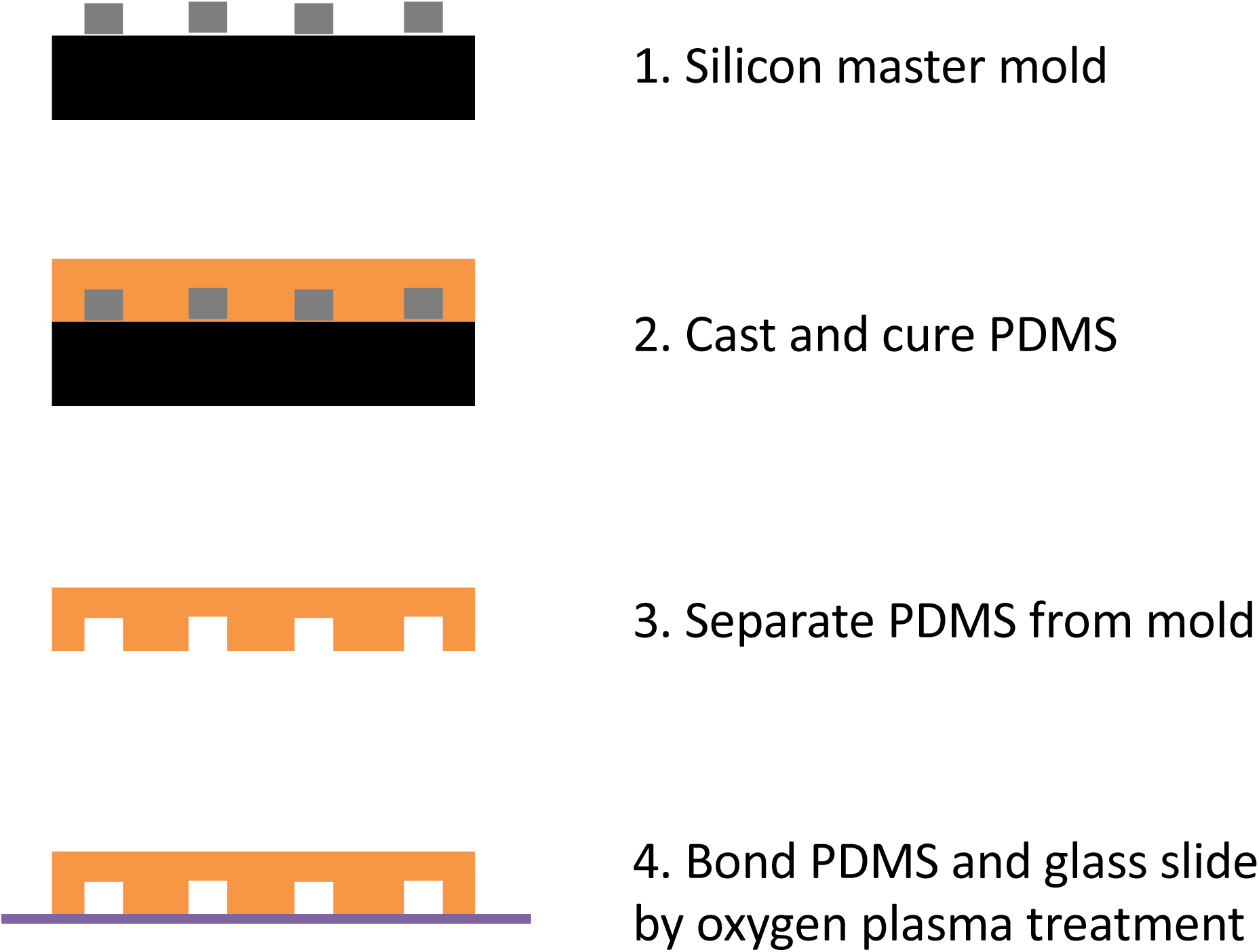
Fabrication of a microfluidic PDMS chip.

**Supplementary Figure 4.**
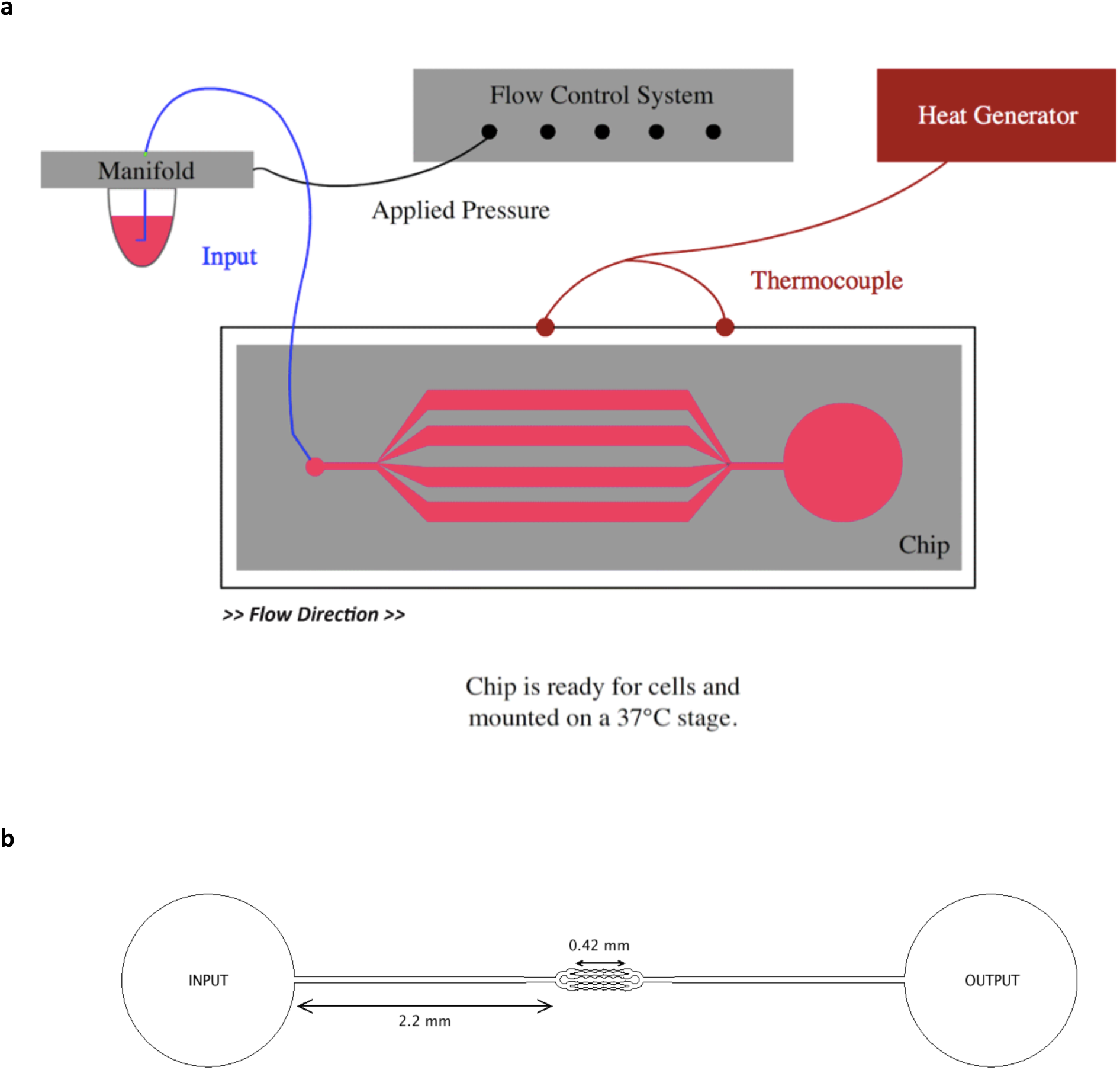
Experimental set-up of the microfluidic system. **(a)** Overall set-up of the microfluidic system. **(b)** The microfluidic chip consists of 4-parallel channels. Each channel is 420 μm in length. The model comes with channels with different lengths of constriction geometry as illustrated in Supplementary Fig. 1.

**Supplementary Table 1.** Summary of transit time data from three types of microfluidic geometry (confined with varying lengths of constrictions) at a constant applied pressure of 10 kPa. Values are presented as (mean ± s.d.). Kruskal-Wallis test. *, significance level at 0.033; **, significance level at 0.002; ***, significance level < 0.001; ns, not significant. **(a)** SK-BR-3 cells. **(b)** MDA-MB-231 cells.

| SK-BR-3 |  |  |  |  |
| --- | --- | --- | --- | --- |
| Parameters | Confined – Type 1<br>(n=73) | Confined – Type 2<br>(n=44) | Confined – Type 3<br>(n=12) | Kruskal-Wallis Test |
| Velocity (mm/s) | 4.86 ± 2.89 | 3.84 ± 3.01 | 2.82 ± 1.91 | P<0.001 (***) |
| Global Transit Time<br>(ms) | 204 ± 339 | 438 ± 822 | 662 ± 1653 | P=0.023 (*) |
| Transit Time across<br>the First Constriction<br>(ms) | 116 ± 253 | 358 ± 769 | 498 ± 1333 | P=0.017 (*) |
| Transit Time (1<sup>st</sup><br>Constriction) / Transit<br>Time (Global) (%) | 41 ± 16 | 54 ± 25 | 55 ± 20 | P=0.013 (*) |

**Supplementary Table 1** Summary of transit time data from three types of microfluidic geometry (confined with varying lengths of constrictions) at a constant applied pressure of 10 kPa.
| MDA-MB-231 |  |  |  |  |
| --- | --- | --- | --- | --- |
| Parameters | Confined – Type 1<br>(n=30) | Confined – Type 2<br>(n=47) | Confined – Type 3<br>(n=38) | Kruskal-Wallis Test |
| Velocity (mm/s) | 7.93 ± 3.54 | 6.36 ± 3.11 | 4.61 ± 2.45 | P<0.001 (***) |
| Global Transit Time<br>(ms) | 73.3 ± 69.3 | 95.5 ± 78.2 | 219 ± 492 | P<0.001 (***) |
| Transit Time across<br>the First Constriction<br>(ms) | 33.7 ± 50.6 | 52.3 ± 71.8 | 172 ± 481 | P<0.001 (***) |
| Transit Time (1<sup>st</sup><br>Constriction) / Transit<br>Time (Global) (%) | 38.9 ± 12.2 | 43.6 ± 17.8 | 55.4 ± 18.9 | P<0.001 (***) |

**Supplementary Table 2.**
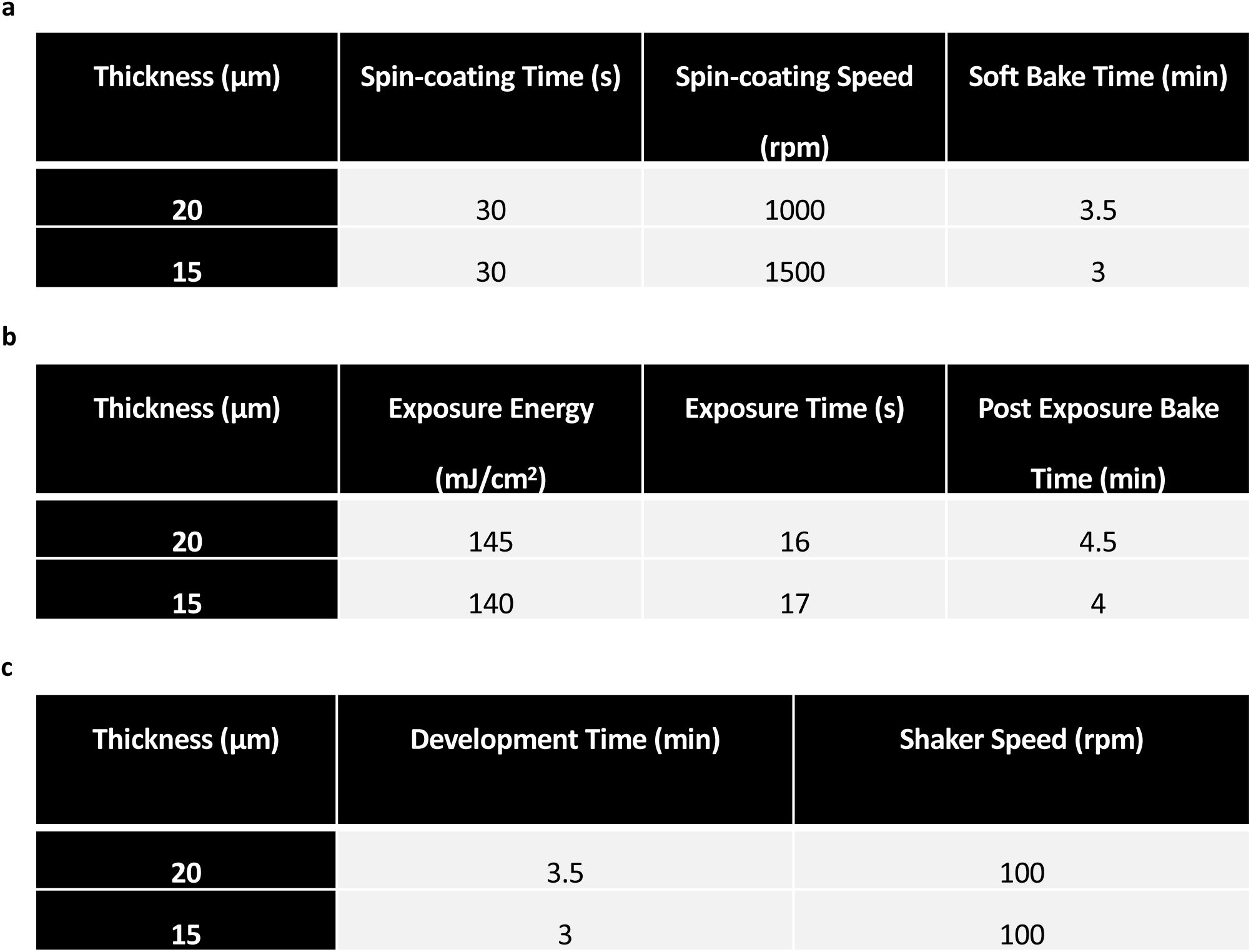
Parameters for optical photolithography. **(a)** Conditions for spin-coating and soft bake. **(b)** Optimal exposure dosage and post exposure bake. **(c)** Development times for SU-8 developer.

**a**
| Thickness ($\mu\text{m}$) | Spin-coating Time (s) | Spin-coating Speed<br>(rpm) | Soft Bake Time (min) |
| --- | --- | --- | --- |
| 20 | 30 | 1000 | 3.5 |
| 15 | 30 | 1500 | 3 |

**b**
| Thickness ($\mu\text{m}$) | Exposure Energy<br>(mJ/cm<sup>2</sup>) | Exposure Time (s) | Post Exposure Bake<br>Time (min) |
| --- | --- | --- | --- |
| 20 | 145 | 16 | 4.5 |
| 15 | 140 | 17 | 4 |

| Thickness ($\mu\text{m}$) | Development Time (min) | Shaker Speed (rpm) |
| --- | --- | --- |
| 20 | 3.5 | 100 |
| 15 | 3 | 100 |

**Supplementary Table 3.**
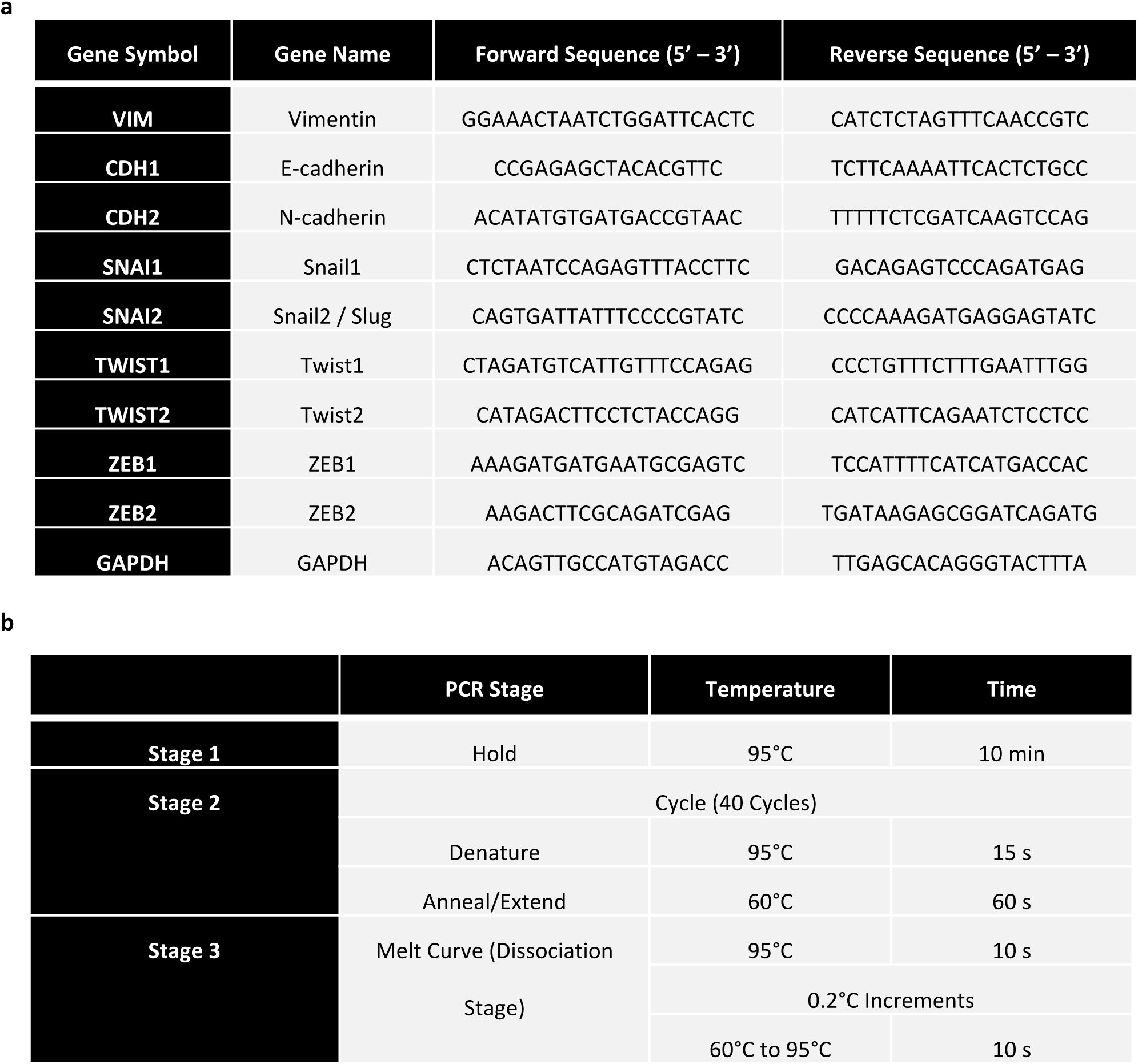
The mRNA primers and thermal cycling conditions for amplification. **(a)** The mRNA primers. **(b)** Thermal cycling conditions for cDNA amplification.

**Supplementary Table 4.**
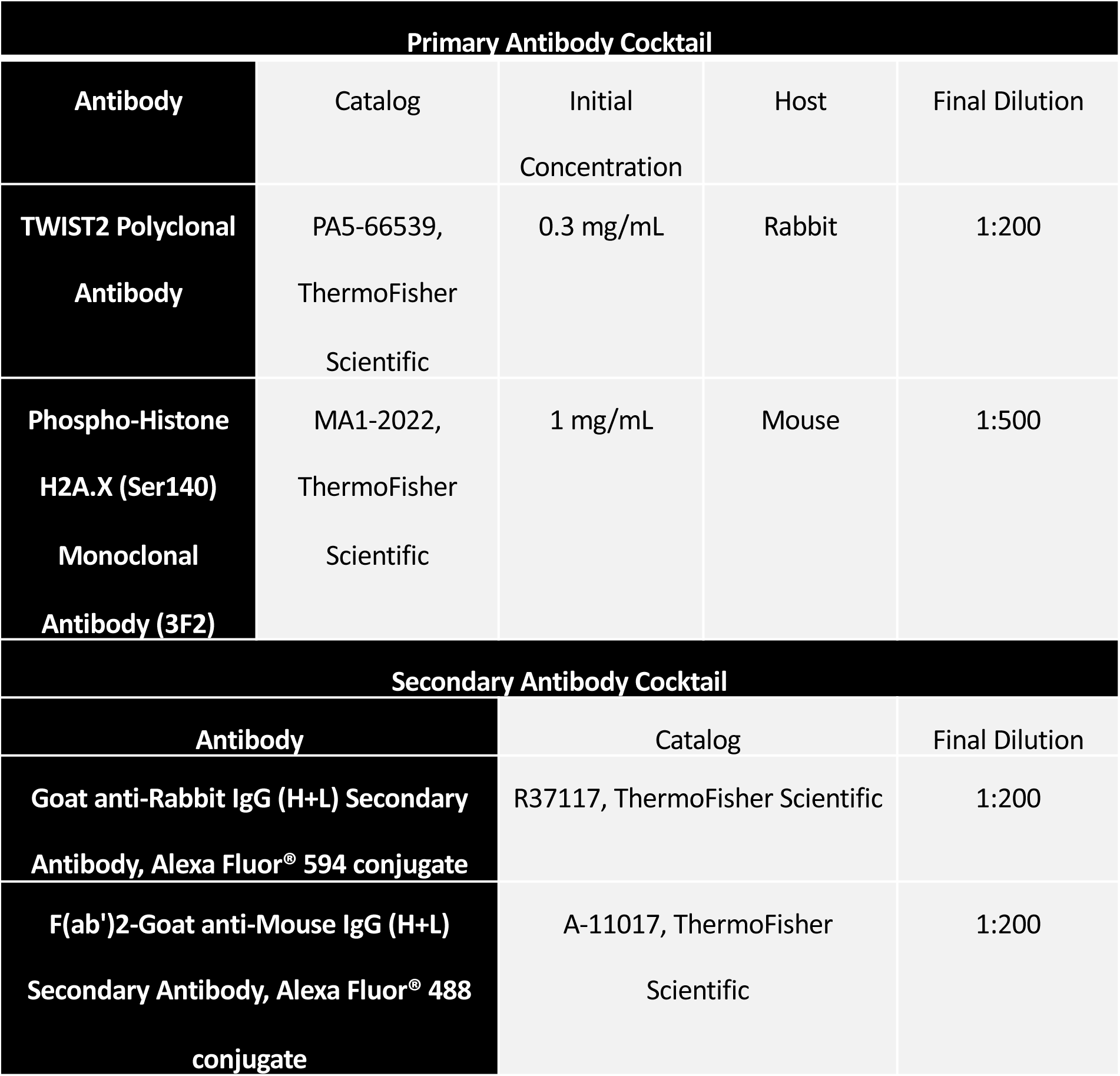
Primary and secondary antibody cocktails.

